# Transition Metal–Weak Organic Acid Synergy Enables Rapid Bactericidal and sporicidal Activity

**DOI:** 10.64898/2026.06.03.729875

**Authors:** Nurit Livnat Levanon, Antara Haldar, Moti Grupper, Oded Lewinson

## Abstract

The global rise in antibiotic-resistant bacteria underscores the urgent need for alternative antimicrobial strategies capable of limiting bacterial growth and viability. We previously demonstrated that weak organic acids and transition metals combine synergistically to strongly inhibit bacterial growth. Here, we extend these findings to determine whether this interaction results only in bacteriostatic suppression or can induce irreversible killing. Using standardized quantitative assays, selected acid–metal combinations achieved rapid bactericidal activity, producing >5-log₁₀ reductions in Gram-negative and Gram-positive bacterial load after brief exposure at concentrations substantially lower than those required for many conventional disinfectants.

This activity extended beyond vegetative cells. *Bacillus cereus* spores, used as a surrogate for spores of *Bacillus anthracis*, exhibited significant loss of viability under defined conditions. Given the environmental persistence of anthrax spores and their relevance in biological threat scenarios, these findings have implications not only for antimicrobial resistance mitigation but also for biodefense and environmental decontamination. Sporicidal efficacy was exposure dependent, with prolonged contact enabling effective killing at reduced concentrations. Together, these results establish acid–metal systems as scalable antimicrobial agents with bactericidal and sporicidal activity relevant to public health and global security.

**Importance statement:** Antibiotic resistance and the persistence of bacterial spores demand new antimicrobial strategies that are both effective and practical. This study shows that combining weak organic acids with transition metals transforms two modest antimicrobial agents into a powerful broad-spectrum killing system. These synergistic formulations rapidly eradicate Gram-negative and Gram-positive bacteria, achieving bactericidal activity at concentrations substantially lower than those used in many conventional disinfectants. Importantly, their activity extends to bacterial spores, among the most resilient biological structures known and a major challenge in environmental decontamination and biodefense. Using *Bacillus cereus* as a surrogate for *Bacillus anthracis*, we demonstrate dramatic sporicidal activity, reducing spore viability by up to six orders of magnitude. By coupling strong efficacy with a simple, scalable, and potentially environmentally compatible formulation, this work establishes transition metal–organic acid combinations as a promising new platform for infection control, environmental decontamination, and biodefense preparedness.

## Introduction

The deliberate release of pathogenic bacteria remains a global security concern, underscoring the need for rapid and reliable decontamination strategies. Among high-risk agents, *Yersinia pestis* and *Bacillus anthracis* represent archetypal biological threats (1–3). *Y. pestis*, the causative agent of plague, can be transmitted via aerosol, leading to rapidly progressing pneumonic disease with high mortality if untreated (4). *B. anthracis*, a spore-forming Gram-positive bacterium, poses a distinct challenge because its endospores are exceptionally resistant to heat, desiccation, radiation, and many chemical disinfectants (5). The resilience and environmental persistence of spores make their eradication particularly demanding in biodefense and environmental decontamination contexts.

While a limited number of chemical disinfectants are available, they often require high concentrations, prolonged exposure times, or harsh formulations, limiting their scalability, environmental compatibility, and suitability for field deployment (5–7). Thus, there is a continued need for antimicrobial strategies that are broad-spectrum, potent, stable, and economically viable.

Transition metals such as copper possess intrinsic antimicrobial activity, largely attributed to intracellular metal toxicity and oxidative damage (8). Weak organic acids are widely used as preservatives and antimicrobial agents in food and agricultural systems (9). Individually, however, both classes typically exhibit moderate potency at concentrations compatible with practical use.

We previously demonstrated that combining weak organic acids with transition metals results in strong synergistic inhibition of bacterial growth (10). The underlying mechanism involves the formation of neutral complexes between metal ions and the deprotonated forms of the organic acids, which enhances membrane permeability by reducing charge and leveraging the hydrophobic character of the acid moiety. This facilitates increased intracellular accumulation of metal ions and amplifies metal-mediated toxicity, producing growth-inhibitory effects across diverse bacterial species. For biodefense applications, while growth inhibition is beneficial, rapid and irreversible killing is by far the preferred outcome. In particular, the ability to eradicate highly resistant spore forms is critical. Whether the organic acid–metal synergy can be extended from growth suppression to robust bactericidal and sporicidal activity remains unknown.

Here, we investigate the bactericidal and sporicidal efficacy of defined organic acid–transition metal combinations against representative Gram-negative and Gram-positive bacteria. *Pseudomonas aeruginosa* and *Staphylococcus aureus* were selected in accordance with international testing standards (BS EN 1040:2005 protocol, (11)), while *Bacillus cereus* and *Yersinia enterocolitica* were included as surrogates for the high-risk pathogens *Bacillus anthracis* and *Yersinia pestis*, respectively (12, 13). Vegetative cells were examined for all species, whereas spore assays were conducted only for *B. cereus*, serving as a model system that mimics the well-established persistence and resilience of anthrax spores in environmental reservoirs.

Using standardized quantitative assays, we distinguish bacteriostatic from bactericidal effects and define the concentration- and exposure-time dependence of killing.

Our findings demonstrate that organic acid–metal combinations can achieve potent bactericidal and sporicidal activity at concentrations substantially lower than those typically required for conventional disinfectants. These results establish a mechanistically grounded and scalable framework for developing environmentally compatible decontamination strategies relevant to biodefense and biosecurity applications.

## Results

### Identification of antibacterial metal–organic acid combinations

To identify effective antibacterial combinations, we performed a systematic matrix screen of six organic acids (acetate, propionate, butyrate, benzoate, formate, sorbate) in combination with three transition metals (Cu²⁺, Zn²⁺, Co²⁺), each tested across three concentrations (Figure 1A). Growth of *Pseudomonas aeruginosa*, *Staphylococcus aureus*, *Bacillus cereus*, and *Yersinia enterocolitica* was monitored in liquid media for 13 hours under standardized conditions (see Methods). These assays provided a rapid and scalable preliminary screen prior to more labor-intensive downstream characterization. Given the large number of experiments (n = 1152), qualitative analysis of the dataset was impractical. We therefore implemented a comparative quantitative framework, inspired by approaches used in high-through-put screening (14). Inspection of representative growth curves revealed a continuum of responses across organisms and conditions (Figure 1B). To facilitate comparison, inhibition was quantified based on endpoint optical density after 800 min relative to untreated controls and discretized into four categories: strong (98–100%) inhibition, partial (90–97%), weak (70–89%), and minimal or no inhibition (<70%).

**Figure 1.**
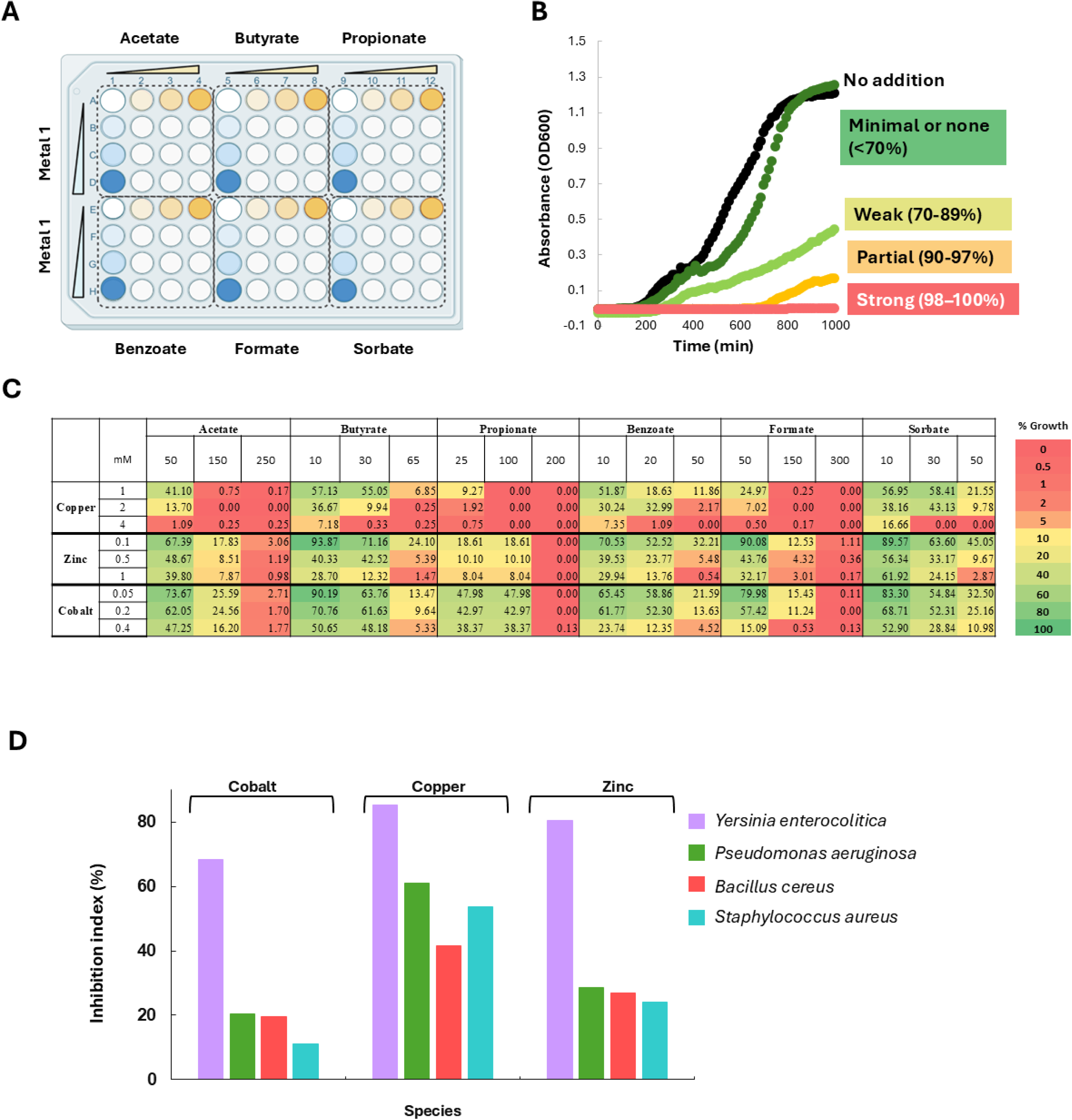
Quantification of inhibitory effects. (A) Representative experimental setup of an experiment used to evaluate combinations of six organic acids (x-axis) with a single metal (y-axis), each at three concentrations, against a single organism. (B) A spectrum of inhibitory effects: *P. aeruginosa* cultures were grown in LB medium for the indicated times, and optical density at 600 nm (OD₆₀₀) was continuously monitored, in the absence (black) or presence of the indicated acid/metal combinations: 50 mM formate/0.1 mM ZnSO₄ (dark green), 20 mM benzoate/0.5 mM ZnSO₄ (light green), 50 mM formate/2 mM CuSO₄ (orange), or 50 mM benzoate/4 mM CuSO₄ (red). Colors match panel C and indicate no, weak, partial, or full inhibition. (C) A heat map summarizing all data for *P. aeruginosa*, with values indicating percent growth relative to growth in the absence of any additives, across all tested combinations of organic acids and metals. Red indicates strong inhibition, orange partial inhibition, yellow weak inhibition, and green no inhibition. (D) Inter-species variability of inhibitory effects. The Inhibition Index was calculated as described in the main text and Methods section, for each metal in combination with the six organic acids across all four tested bacteria. As shown, *Yersinia enterocolitica* emerged as the most susceptible, *Staphylococcus aureus* as the least, while *Pseudomonas aeruginosa* and *Bacillus cereus* displayed intermediate susceptibility.

These classifications were used to generate heat maps summarizing the dataset (Figure 1C; Supplementary Figure 1) and to derive an inhibition index, enabling comparison of treatment efficacy across conditions. Using the formulation of Zhang et al (14), the inhibition index was calculated as a weighted fraction of conditions exhibiting inhibitory effects, with strong and partial inhibition assigned weights of 1 and 0.5, respectively, while weaker effects were excluded:

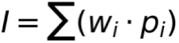

Where *wi* denotes the assigned weight for each inhibition category and *pi* the fraction of conditions in that category (see method for details). This metric therefore captures the prevalence of robust inhibitory responses across the tested conditions.

This approach enabled rapid identification of active and inactive regions of the parameter space. Heat maps revealed clear organism-specific patterns, allowing prioritization of conditions that consistently produced strong inhibition (Figure 1C, Supplementary Figure 1). Notably, no combination was universally effective; instead, responses were highly organism-dependent. For example, acetate–copper combinations were highly effective against *Staphylococcus aureus* but largely ineffective against *Pseudomonas aeruginosa*, whereas sorbate–copper combinations showed activity against *S. aureus* and were markedly more effective against *P. aeruginosa*.

Consistent with this variability, the inhibition index revealed differences in overall susceptibility. *Yersinia enterocolitica* was the most susceptible, followed by *Pseudomonas aeruginosa*, while *Staphylococcus aureus* and *Bacillus cereus* were more resistant (Figure 1D). Within this limited set, this trend aligns with Gram classification, with Gram-negative species showing greater sensitivity than Gram-positive species.

### Bactericidal activity of mixtures of organic acids and transition metals

The above results (Figure 1, Supplementary Figure 1) demonstrate strong growth inhibition by selected metal–organic acid combinations; however, they do not distinguish between growth suppression and irreversible cell death (i.e., bacteriostatic versus bactericidal effects).

Bactericidal activity was evaluated according to the BS EN 1040:2005 quantitative suspension protocol, which defines bactericidal efficacy as a ≥5-log₁₀ reduction in viable bacteria under standardized conditions (see methods for full details).

Briefly, exponentially growing cultures (∼4 × 10⁸ CFU/mL) were exposed to the indicated metal–acid mixtures for 5 minutes at 20°C. Reactions were then neutralized by dilution, and serial dilutions were plated on TSA plates to enumerate surviving colony-forming units (CFU). Water-treated samples served as controls. A ≥5-log₁₀ reduction in CFU relative to control was considered bactericidal.

Comparison of these CFU-based killing results with the prior OD-based growth inhibition assays revealed a clear divergence between growth suppression and irreversible cell death. Several combinations that strongly inhibited growth did not reduce the number of viable colonies indicating bacteriostatic rather than bactericidal activity under those conditions. For example, several combinations of benzoate with either copper or zinc completely abolished growth (Supplementary Figure 1C) but had no bactericidal activity (Supplementary Figure 2). Importantly, for each of the tested organisms, we identified combinations that produced rapid and irreversible killing and reduced the CFU count by at least 10^7^ (Figure 2).

**Figure 2.**
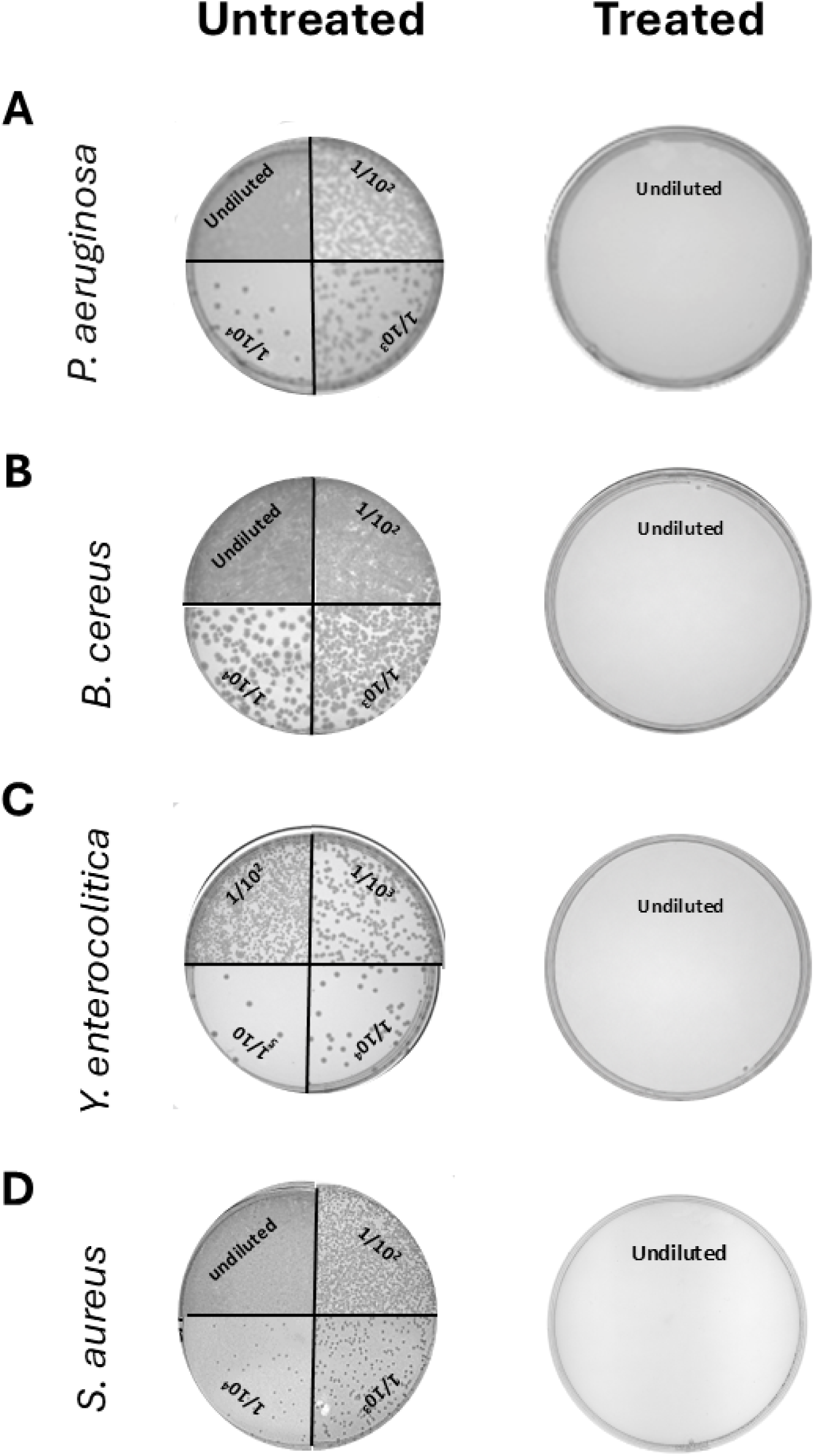
Bactericidal activity of mixtures of organic acids and transition metals. Exponentially growing cultures (∼4 × 10⁸ CFU/mL) of the indicated bacteria were exposed for 5 min to H₂O (left panels, untreated) or to (A) 2% acetic acid/2 mM CuSO₄, (B) 0.2% formic acid/0.5 mM CuSO₄, (C) 0.4% formic acid/1 mM CuSO₄, or (D) 1% formic acid/2.5 mM CuSO₄. Serial dilutions (as indicated) were plated for CFU enumeration. Solid black lines mark boundaries between images assembled from plates of different dilutions. For treated samples, only undiluted plates are shown, as all other dilutions similarly yielded no CFUs. Images are representative of at least three independent experiments.

### Determination of Minimum Bactericidal Concentrations (MBC) reveals interspecies variability in bactericidal efficacy

Next, for each species, we selected an effective transition metal–organic acid combination and quantified its inhibitory potency by determining the Minimum Bactericidal Concentration (MBC), defined as the lowest concentration of the mixture that completely abolished detectable bacterial growth.

Interestingly, in all cases tested, the dose–response relationship showed a sharp transition from no effect to near-complete killing, consistent with a threshold-like bactericidal response (Figure 3).

**Figure 3.**
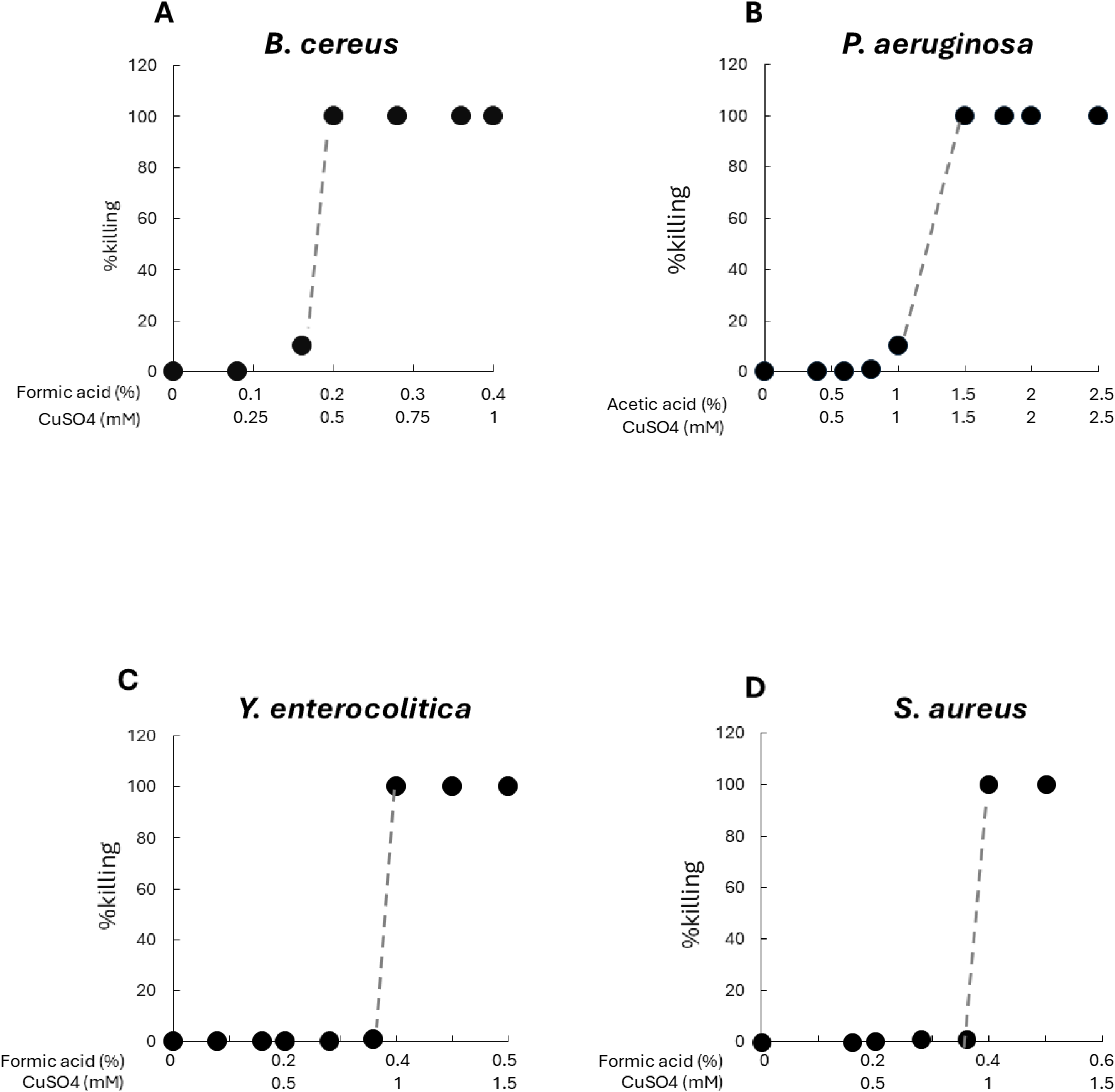
Determination of Minimum Bactericidal Concentrations (MBC) Dose–response relationships showing percentage killing as a function of increasing concentrations of organic acid–copper mixtures (as indicated) for Bacillus cereus (A), Pseudomonas aeruginosa (B), Yersinia enterocolitica (C), and Staphylococcus aureus (D). Each point represents the percentage reduction in colony-forming units (CFU) relative to the untreated control. Dashed lines indicate the sharp transition from minimal or no killing to complete killing.

Bactericidal thresholds varied markedly across species. *Bacillus cereus* emerged as the most susceptible organism, with 0.2% formic acid combined with 0.5 mM CuSO₄ producing a >5-log₁₀ reduction after 5 minutes of exposure. *Yersinia enterocolitica* and *Staphylococcus aureus* exhibited intermediate sensitivity, requiring approximately twofold higher concentrations (0.4% formic acid and 1 mM CuSO₄) to achieve complete bactericidal activity. *Pseudomonas aeruginosa*, the most resistant organism tested, required higher concentrations (1.5% acetic acid + 1.5 mM CuSO₄) for eradication.

Notably, this sensitivity hierarchy differs from that observed in the growth inhibition assays (Figure 1D), where *Yersinia enterocolitica* appeared the most sensitive, while *Bacillus cereus* showed comparatively lower susceptibility, providing an additional demonstration of the distinction between bacteriostatic and bactericidal effects.

Given that the species-specific bactericidal thresholds were defined under 5-minute exposure conditions, we next asked whether similar killing efficacy could be achieved at lower concentrations by extending the duration of exposure. To address this, we performed time-dependent killing experiments using sub-bactericidal concentrations. These analyses revealed that bactericidal activity was strongly exposure-duration dependent, with progressive reductions in CFU observed over extended incubation periods (Supplementary Figures 3-5). In several cases, prolonged contact partially or fully compensated for reduced concentration, indicating that concentration and time function as interdependent parameters in determining killing efficacy.

### Sporicidal activity of transition metal–weak organic acids combinations

Having established potent bactericidal activity against vegetative cells, we next asked whether this approach could overcome the far greater challenge posed by bacterial endospores. Sporicidal activity remains difficult to achieve in both clinical and biodefense settings, as only a limited number of chemical agents are capable of reliably inactivating spores, often requiring high concentrations, prolonged exposure, or harsh conditions that limit environmental compatibility. The need for effective and scalable sporicidal strategies is therefore pressing in the contexts of infection control, environmental decontamination, and biological threat preparedness.

Among spore-forming pathogens, *Bacillus anthracis* is widely recognized as a high-priority biodefense threat due to the extreme environmental persistence and resilience of its spores. Given the regulatory and biosafety constraints associated with working with *B. anthracis*, we employed *Bacillus cereus* as a surrogate organism, which is commonly used in biodefense and decontamination research. The two species share >99% chromosomal sequence identity and exhibit highly conserved spore architecture and intrinsic spore resistance mechanisms.

A purified preparation of *Bacillus cereus* spores was obtained essentially as previously described (15). Characteristic endospore structures were clearly visible after 5 days of growth on nutrient agar supplemented with 100 μM MnCl₂, as visualized by malachite green–safranin staining (Figure 4). Treatment with 50% ethanol (1 h, room temperature), which completely eliminated vegetative cells (Figure 4A,B), did not reduce CFU counts in the spore preparations (Figure 4C,D), demonstrating that colony formation arises from ethanol-resistant spores and that vegetative cells are absent or below the limit of detection. Together, these results demonstrate that the preparation is highly purified and suitable for quantitative analysis.

**Figure 4.**
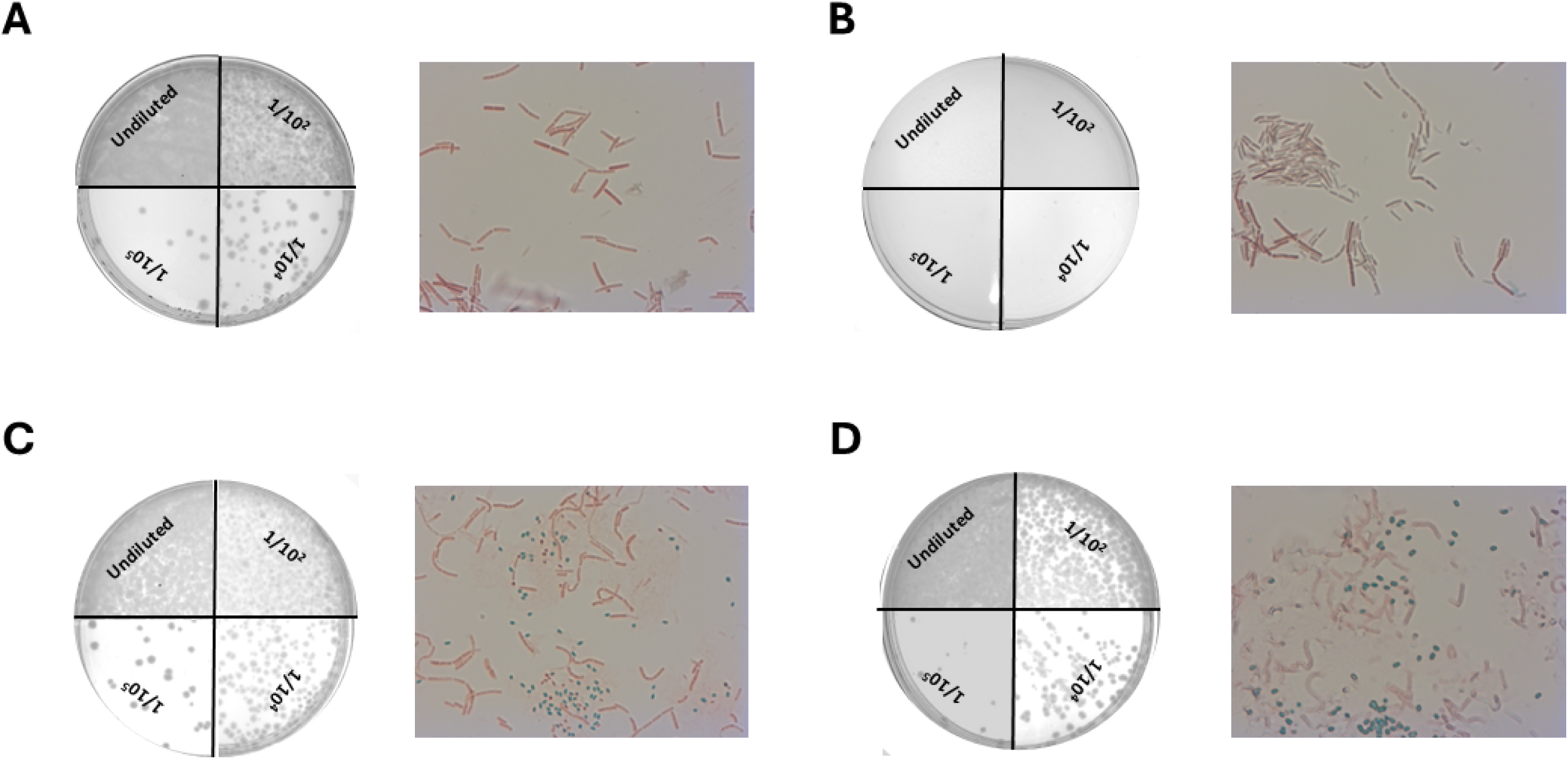
Preparation and purification of *Bacillus cereus* spores. Vegetative cells (A,B) or purified spores (C,D) of *Bacillus cereus* were plated on TSA plates before (A,C) or after (B,D) treatment with 50% ethanol (1h, room temperature). The corresponding images show malachite green/safranin staining, visualized at ×63 magnification under oil immersion. Images are representative of at least three independent experiments.

We first examined whether metal–acid mixtures could prevent spore outgrowth. To this end, spores were prepared as described above, and residual vegetative cells were eliminated by treatment with 50% ethanol. We then tested concentrations that were bactericidal for vegetative cells (Figure 3). These conditions completely abolished spore outgrowth in both liquid (Figure 5A) and solid (Figure 5B–D) media.

**Figure 5.**
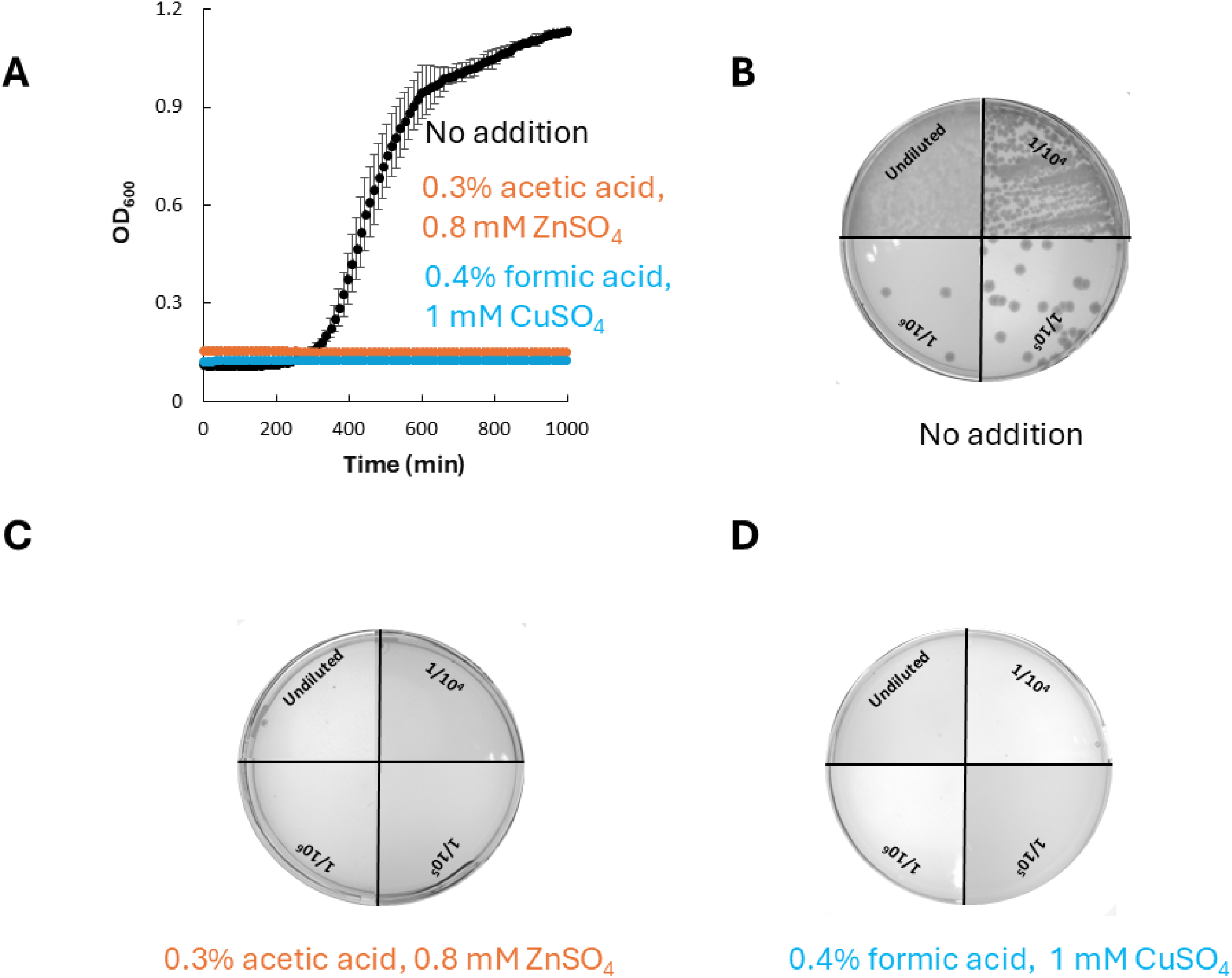
Inhibition of outgrowth of *Bacillus cereus* spores. (A) Spores of *Bacillus cereus* were prepared as described in the Methods section and cultured in LB medium in the absence or presence of the indicated combinations of weak organic acids and transition metals. Error bars (shown unless smaller than the symbols) represent standard deviations of triplicate measurements. (B–D) Spores were serially diluted (as indicated) and plated on TSA plates (B) or TSA plates supplemented with the indicated combinations of weak organic acids and transition metals (C, D). Solid black lines mark boundaries between images assembled from plates of different dilutions. Images are representative of at least three independent experiments.

To determine whether the observed inhibition reflected loss of viability rather than delayed germination, we performed quantitative spore killing assays. Purified spore preparations were exposed to the indicated metal–acid mixtures for 1 h at 20 °C, after which the spores were washed in DDW and plated on solid medium for CFU enumeration.

Unsurprisingly, spores exhibited markedly greater resistance than vegetative cells. While vegetative cells of *Bacillus cereus* were efficiently killed by as little as 0.15% acetic acid combined with 0.4 mM ZnSO₄, spores were only minimally affected even by ∼100-fold higher concentrations (Figure 6A–C). Notably, high concentrations of Cu/formic acid (20% formic acid + 50 mM CuSO₄) reduced viable spore counts by approximately 5–6 log₁₀ (Figure 6D), highlighting the potential of this combination as an effective sporicidal agent.

**Figure 6.**
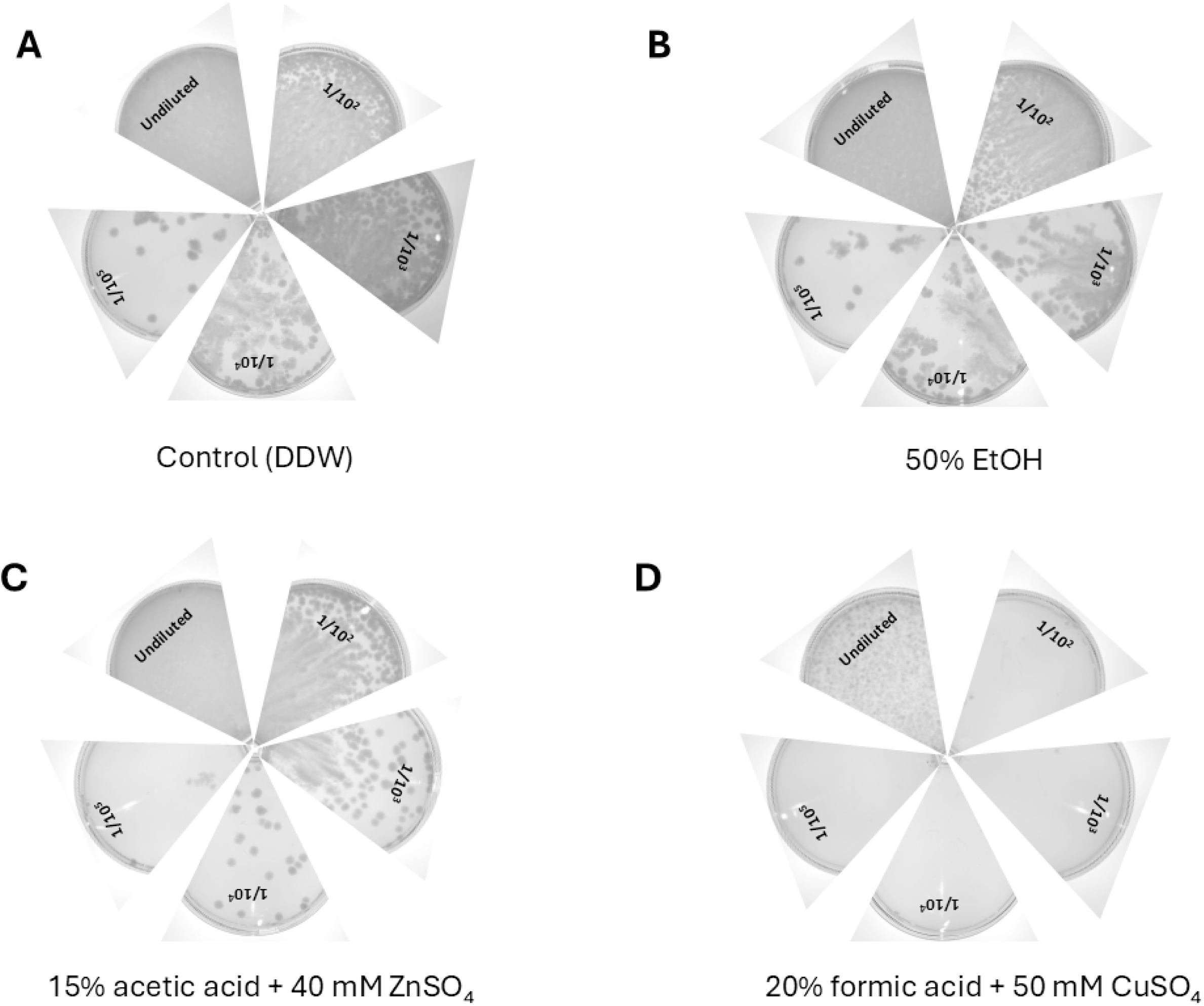
Sporicidal activity of combinations of weak organic acids and transition metals. Spores of *Bacillus cereus* were prepared as described in the Methods section and exposed for 3 h at 20 °C to H₂O (A), 50% ethanol (B), 15% acetic acid/40 mM ZnSO₄ (C), or 20% formic acid/50 mM CuSO₄ (D). Spores were then washed, serially diluted (as indicated), and plated on TSA plates. The formic acid/CuSO₄ combination reduced spore CFU counts by 10³–10⁴ (D). Images are representative of at least three independent experiments.

## Discussion

In this study, we demonstrate that combinations of weak organic acids and transition metals can be translated from synergistic growth inhibition into rapid and irreversible bactericidal activity across diverse Gram-negative and Gram-positive bacteria. While previous work established strong inhibitory synergy between organic acids and transition metals, the present data extend those findings by systematically distinguishing growth suppression from true loss of viability and by quantifying bactericidal thresholds under standardized conditions.

### From inhibition to killing

High-throughput liquid assays identified multiple metal–acid combinations that completely suppressed optical density increase. However, OD-based measurements do not differentiate between bacteriostatic and bactericidal effects. By applying plate-based killing assays according to the BS EN 1040:2005 framework, we demonstrate that only a subset of these inhibitory combinations achieve ≥5-log₁₀ reduction within 5 minutes. This divergence highlights an important conceptual distinction: complete growth inhibition does not necessarily imply irreversible cell death.

### Species-dependent susceptibility

Marked differences in bactericidal thresholds were observed across species. Overall, the two Gram-positive organisms exhibited greater resistance than their Gram-negative counterparts. We do not attribute this difference to the presence of more elaborate copper defense or metal efflux systems, which are generally more extensive in Gram-negative bacteria. Rather, it likely reflects differences in cell envelope architecture, with the thicker peptidoglycan-rich envelope of Gram-positive bacteria providing a more effective barrier to diffusion and intracellular accumulation of toxic metal–acid complexes.

Importantly, bactericidal activity was achieved at concentrations substantially lower than those commonly used for conventional disinfectants, indicating favorable potency relative to currently available decontamination agents.

### Exposure time as a determinant of efficacy

A key finding of this study is the strong exposure-time dependence of killing. In these experiments (Supplementary Figure 3-5), we used sublethal concentrations below the minimal bactericidal concentrations. Under these conditions, increasing the incubation time dramatically improved killing efficacy, in some cases by several orders of magnitude. This result has practical implications. In real-world decontamination scenarios, disinfectants applied to contaminated surfaces are likely to remain in contact for extended periods. The evaporation of organic acids and gradual dilution of residual metal salts suggest that sustained, low-level exposure may contribute significantly to antimicrobial efficacy. Thus, concentration and time can be viewed as interchangeable parameters within defined limits.

### Activity against spores

The extension of these analyses to purified endospores further clarifies both the strengths and limitations of the system. As expected, spores exhibited markedly greater resistance than vegetative cells. Notably, concentrations that were bactericidal against vegetative cells were also sufficient to effectively inhibit spore germination and outgrowth. In contrast, direct sporicidal activity required substantially higher concentrations, approximately two orders of magnitude greater than those effective against vegetative cells. These findings underscore the fundamental structural and physiological resilience of bacterial endospores. The multilayered spore coat, low metabolic activity, and restricted permeability likely limit intracellular metal accumulation and oxidative damage.

### Mechanistic considerations

We previously demonstrated that combining weak organic acids with transition metals results in strong synergistic inhibition of bacterial growth (10). Using both liposomes and whole cells, we showed that weak organic acids enhance the membrane permeability of transition metals, leading to increased intracellular metal accumulation and amplified metal-mediated toxicity across diverse bacterial species. This effect may result either from the formation of neutral metal–organic acid complexes, whose reduced charge facilitates membrane permeation, or from organic acid–mediated destabilization of the membrane, which enhances metal entry.

The present study may lend support to the latter mechanism. In all species tested, the bactericidal dose–response curves exhibited a remarkably sharp transition from little or no killing to complete killing. Such threshold-like behavior is difficult to reconcile with a simple dependence on bulk metal concentration and instead suggests that bactericidal activity may depend on reaching a critical local concentration at the membrane surface, consistent with organic acid–mediated membrane destabilization facilitating metal entry.

Future studies will be required to distinguish between these and additional possible mechanisms, and to determine whether vegetative cell killing and sporicidal activity occur through distinct mechanisms.

### Implications for biodefense and decontamination

While the present study demonstrates clear sporicidal activity, achieving high-level spore killing currently requires relatively high concentrations. Further optimization will therefore be needed to obtain efficient sporicidal efficacy at lower concentrations. Such optimization may be achievable through the addition of compounds that increase local surface concentration or partially destabilize the spore coat (16–18), or by modifying the stoichiometric ratio between the organic acid and the transition metal. Nevertheless, the broad-spectrum activity, rapid bactericidal action, exposure-dependent flexibility, and meaningful activity against spores demonstrated here support the feasibility of weak organic acid–transition metal systems as scalable and environmentally compatible decontamination agents and provide a strong foundation for future development.

## Materials and Methods

### Bacterial strains and growth conditions

*Pseudomonas aeruginosa* ATCC 15442, *Staphylococcus aureus* ATCC 6538, *Bacillus cereus* ATCC 11778, and *Yersinia enterocolitica* ATCC 9610 were used throughout this study. Unless otherwise indicated, cultures were grown in LB or Tryptic Soy Broth with constant shaking. *P. aeruginosa* and *S. aureus* were cultured at 37 °C, whereas *B. cereus* and *Y. enterocolitica* were cultured at 30 °C. Overnight cultures were diluted into fresh medium to the indicated optical densities prior to each experiment.

### Preparation of organic acid–metal mixtures

Stock solutions of organic acids and transition metal salts were prepared in sterile double distilled water. Organic acids tested included the sodium salts of acetate, butyrate, propionate, benzoate, formate, and sorbate. Transition metals were supplied as CuSO₄, ZnSO₄, or CoSO₄.

A master plate with the combinations at 2X concentrations was set (one master plate resulting combinations of one metal salt with 6 acid anions). Master plates were stored at 4°C, protected from light.

### Growth assays

Growth assays were performed in 96-well plates using combinations of weak organic acids and transition metal salts at the indicated concentrations. Bacterial cultures were diluted to an optical density at 600 nm of 0.01 in LB. A volume of 75 µL culture was dispensed into each well containing the 75 uL of the 2x master plate resulting indicated concentrations of organic acids and/or metal salts.

Plates were incubated in an automated plate reader (Infinite M200 Pro, Tecan) with intermittent shaking at 37 °C (PSA and SA) or 30 °C (BC or YE). Optical density at 600 nm (OD600) was recorded at regular intervals for the indicated durations.

### Heat map generation and calculation of inhibition index

Growth inhibition was quantified using endpoint optical density measurements obtained after 800 min of incubation. For each condition, OD600 values were normalized relative to untreated controls and expressed as percentage growth relative to untreated samples. Based on the normalized growth values, inhibitory effects were classified into four categories: strong (98–100% inhibition), partial (90–97%), weak (70–89%), and minimal or no inhibition (<70%). Heat maps summarizing inhibition profiles across all tested combinations were generated using Microsoft Excel conditional formatting based on the categorized inhibition datasets.

To enable quantitative comparison across conditions, a semi-quantitative inhibition index was calculated using a weighted scoring approach based on the distribution of categorical responses across the heat maps (14, 19). The inhibition index was calculated according to the following equation:

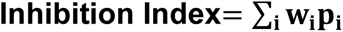

where *w*_*i*_ represents the weight assigned to each inhibition category and *p*_*i*_represents the fraction of conditions belonging to that category. Red and orange categories were assigned weights of 1 and 0.5, respectively, whereas yellow and green categories were assigned a weight of 0.

Under these conditions, the equation simplifies to:

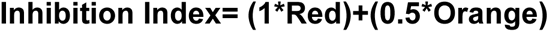

This approach provides a semi-quantitative summary of inhibitory efficacy while preserving the categorical structure of the dataset.

### Determination of bactericidal activity

Assays are based on the European standard protocol for evaluation the bactericidal activity of chemical disinfectants and antiseptics (11). Briefly, exponentially growing cultures (∼10⁸ CFU mL⁻¹) were exposed to the indicated combinations of organic acids and transition metals for 5 min at 20°C. Following exposure, samples were immediately diluted 10-fold in fresh medium to terminate the treatment.

Samples were then serially diluted in water and plated on Tryptic Soy Agar plates. Plates were incubated for colony formation (overnight at 37 °C unless otherwise indicated). Colony-forming units (CFUs) were enumerated, and survival was calculated relative to untreated control samples. Plates were documented using the Vilber Fusion FX7 Edge Spectra.

### Determination of Minimum Bactericidal Concentrations (MBC)

Minimum Bactericidal Concentrations (MBC) values were determined using plate-based killing assays. Bacterial cultures were exposed to decreasing concentrations of selected organic acid–metal mixtures for 5 min at 20 °C. Following exposure, compounds were removed by dilution in fresh medium. Cells were serially diluted and plated on TSA plates. The Minimum Bactericidal Concentration (MBC) was defined as the lowest concentration resulting in complete loss of colony formation.

### Time-dependent killing assays

Time-dependent killing experiments were performed by exposing bacterial cultures to selected organic acid–metal mixtures for varying exposure times ranging from several minutes to several hours. Following treatment, samples were diluted to remove residual compounds, serially diluted, and plated on TSA plates.

Colonies were enumerated after incubation, and bacterial survival was calculated as a function of exposure time relative to untreated controls.

### Preparation of *Bacillus cereus* spores

Spores of *Bacillus cereus* ATCC 11778 were prepared under nutrient-limiting conditions as previously described (15). Several colonies from rich media agar plate (TSA or LB) were grown overnight at 30 °C in double-strength Schaeffer’s-glucose (2×SG) medium containing nutrient broth (Foremedium 16g/L)), KCl (2 g/L), MgSO₄·7H₂O (0.5g/L),1 mM Ca(NO₃)₂, 100 μM MnCl₂ (supplier name),1 μM FeSO₄, and 0.1 % glucose.

Cultures were then spread on nutrient agar plates supplemented with 100 µM MnCl₂ and incubated at 30 °C for five days to induce sporulation.

To eliminate residual vegetative cells, spore preparations were treated with 50% ethanol for 1 h at room temperature. Sporulation efficiency was verified by ethanol resistance assays and by malachite green–safranin staining.

### Malachite green strain

Bacterial spores were stained using the Schaeffer–Fulton method (20) with malachite green as the primary stain. A thin smear of the bacterial culture was first prepared on a clean glass microscope slide using a small volume sores’ suspension. The smear was air-dried and then heat-fixed by briefly passing the slide through a flame. The slide was placed over a steaming water bath, and a piece of absorbent paper was positioned over the smear. The smear was flooded with 5% (w/v) malachite green solution and steamed for 5 minutes, ensuring the stain remained moist throughout the process by adding additional stain as needed. Following staining, the slide was allowed to cool and then rinsed gently with DDW to remove excess dye. This step decolorizes vegetative cells while retaining stain within spores.

The smear was then counterstained with safranin for 2 minutes, rinsed again with water, and gently blotted dry with absorbent paper.

Prepared slides were examined under a light microscope at 63x magnification using oil immersion. Endospores appeared green, while vegetative cells-stained red.

### Spore outgrowth inhibition assays

*Bacillus cereus* ATCC 11778 spores were prepared as described above. Outgrowth inhibition was done in liquid or solid rich media. For the first, spores’ preparation was diluted (96-well plates) and final volume of 150 μl with the indicated concentrations of organic acids and transition metals. For outgrowth inhibition on agar plates, spores were resuspended in double-distilled water (DDW) and serially diluted (undiluted, 1:100, 1:1,000, 1:10,000, and 1:100,000). A volume of 250 µL from each dilution was plated on Tryptic Soy Agar (TSA) plates. Following plating, spores were allowed to settle on rich medium, and initiate germination for 2 h at room temperature. Subsequently, 0.2 mL of treatment solutions was evenly spread on the surface of each plate using a sterile spreader, yielding the indicated final concentrations within the agar.

Plates were incubated at 30 °C overnight to allow colony formation. Colony-forming units (CFUs) were enumerated, and inhibition of spore outgrowth was assessed relative to untreated controls.

### Spore killing assays

*Bacillus cereus* ATCC 11778 spores were prepared as described above. Prior to every experiment, spores were harvested from nutrient agar plates supplemented with 100 µM MnCl₂ and resuspended in double-distilled water (DDW). To eliminate residual vegetative cells, spore suspensions were treated with 50% ethanol for 1 h at room temperature, followed by centrifugation (15 min, 3,000×g) and two washes with DDW.

For killing assays, spore suspensions were exposed to the indicated treatments at 20 °C. Following treatment, samples were centrifuged (15 minutes, 3,000×g) and resuspended in 1 mL DDW to remove residual compounds. Samples were then serially diluted in DDW (undiluted, 1:100, 1:1,000, 1:10,000, and 1:100,000), and 500 µL of each dilution was plated on Tryptic Soy Agar (TSA) plates. Plates were incubated overnight at 30 °C, and colony-forming units (CFUs) were enumerated to assess spore survival relative to untreated controls.

## Data analysis

All experiments were performed with at least three independent biological replicates, each containing technical duplicates or triplicates unless otherwise stated. Data were analyzed using standard statistical methods and results are presented as means of independent experiments.

## Funding

This research was supported by the Ministry of Defense (Grants No. 44441253057 and 4441399304)

## Conflict of Interests

The authors declare no conflict of interest (such as defined by ASM policy).

## Author contribution

Conceptualization: O.L., N.L.L. Methodology: A.H., M.G., N.L.L., O.L.

Investigation: A.H., M.G., N.L.L., O.L., Analysis: A.H., M.G., N.L.L., O.L.

Writing, review and editing: A.H., M.G., N.L.L., O.L.

---

**Supplementary Figure 1.**
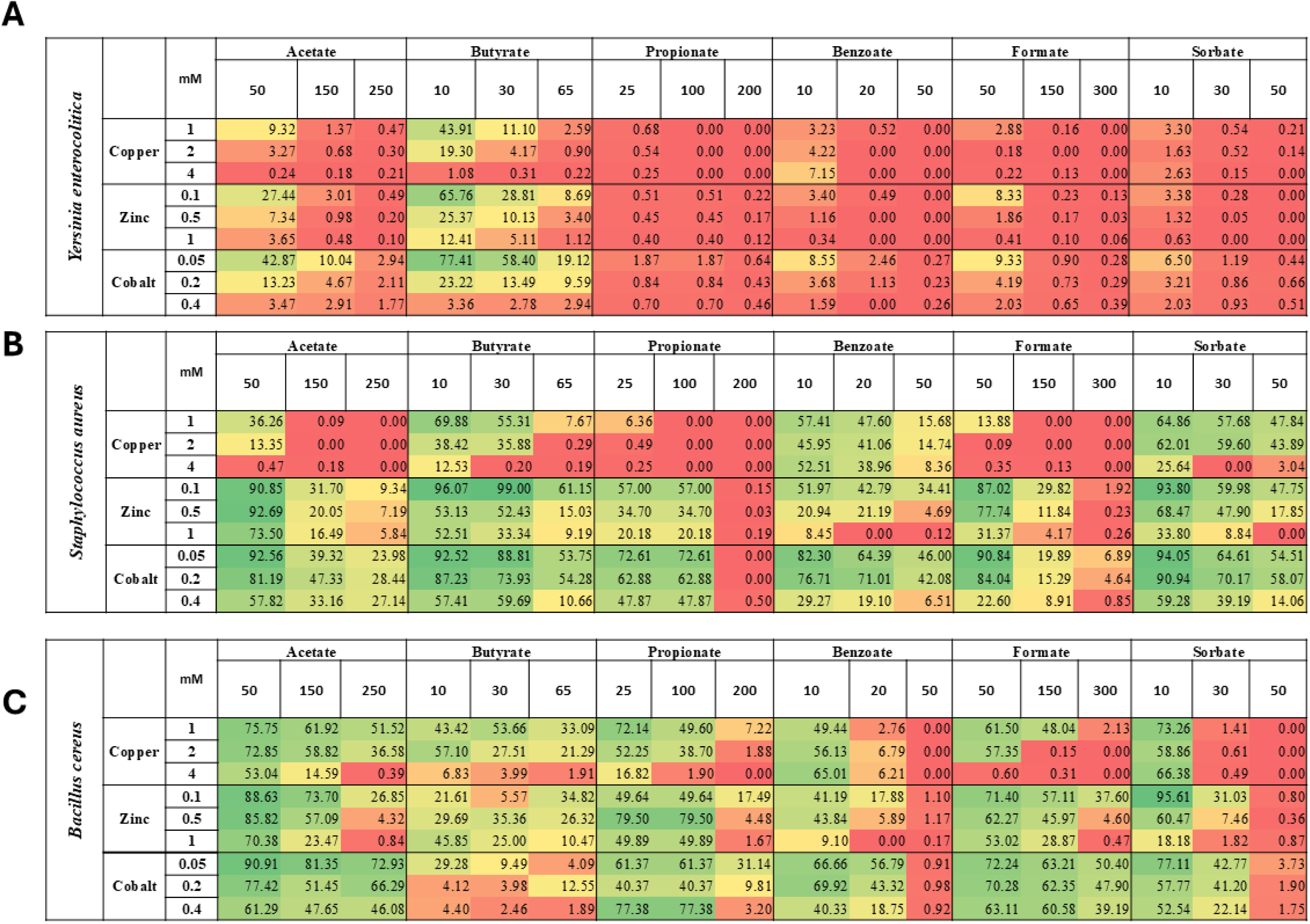
Interspecies variability in treatment efficacy. Heat maps summarizing all data for *Yersinia enterocolitica, Staphylococcus aureus,* and *Bacillus cereus,* as indicated. The values indicate percent growth relative to growth in the absence of any additives, across all tested combinations of organic acids and metals. Red indicates strong inhibition, orange partial inhibition, yellow weak inhibition, and green no inhibition.

**Supplementary Figure 2.**
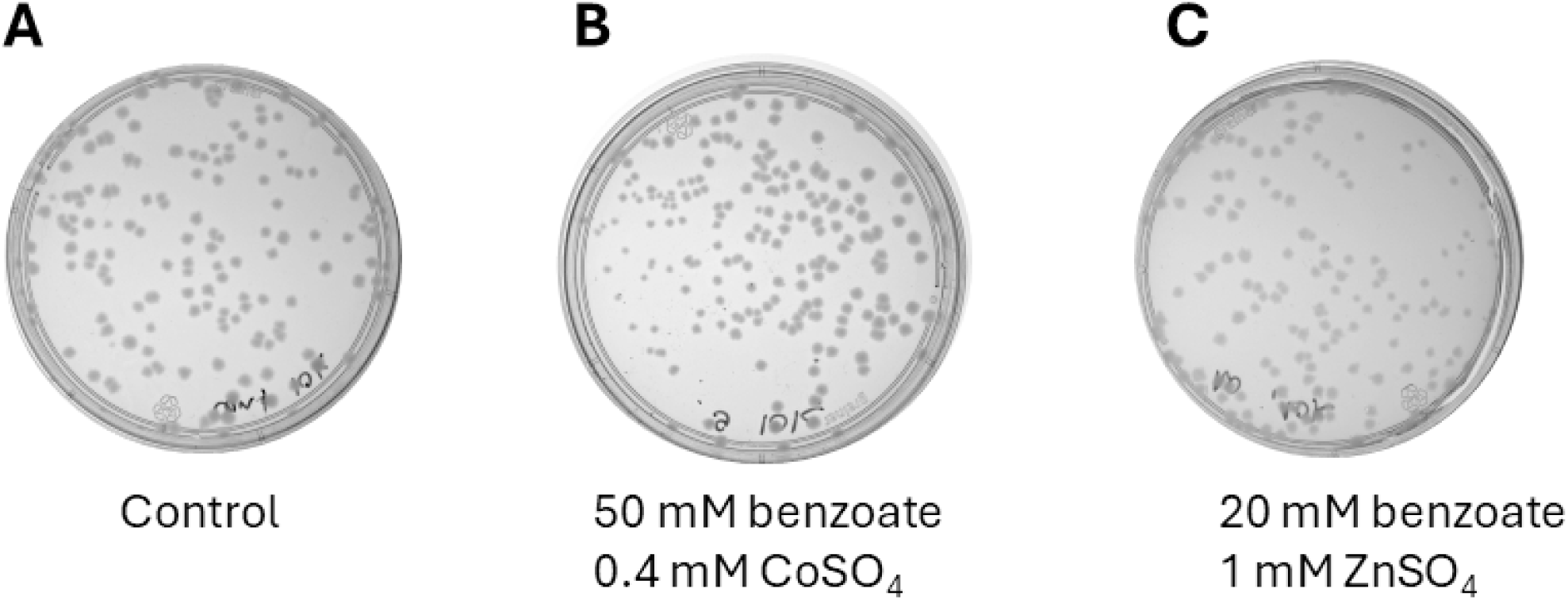
No bactericidal activity for benzoate mixtures that inhibited growth. Exponentially growing cultures (∼4 × 10⁸ CFU/mL) of *B.cereus* were exposed for 5 min to H₂O (A) or to (B) 50 mM benzoate/0.4 mM CoSO₄, (C) 20 mM benzoate/1 mM ZnSO4₄. Shown is 1/10^4^ dilution.

**Supplementary Figure 3.**
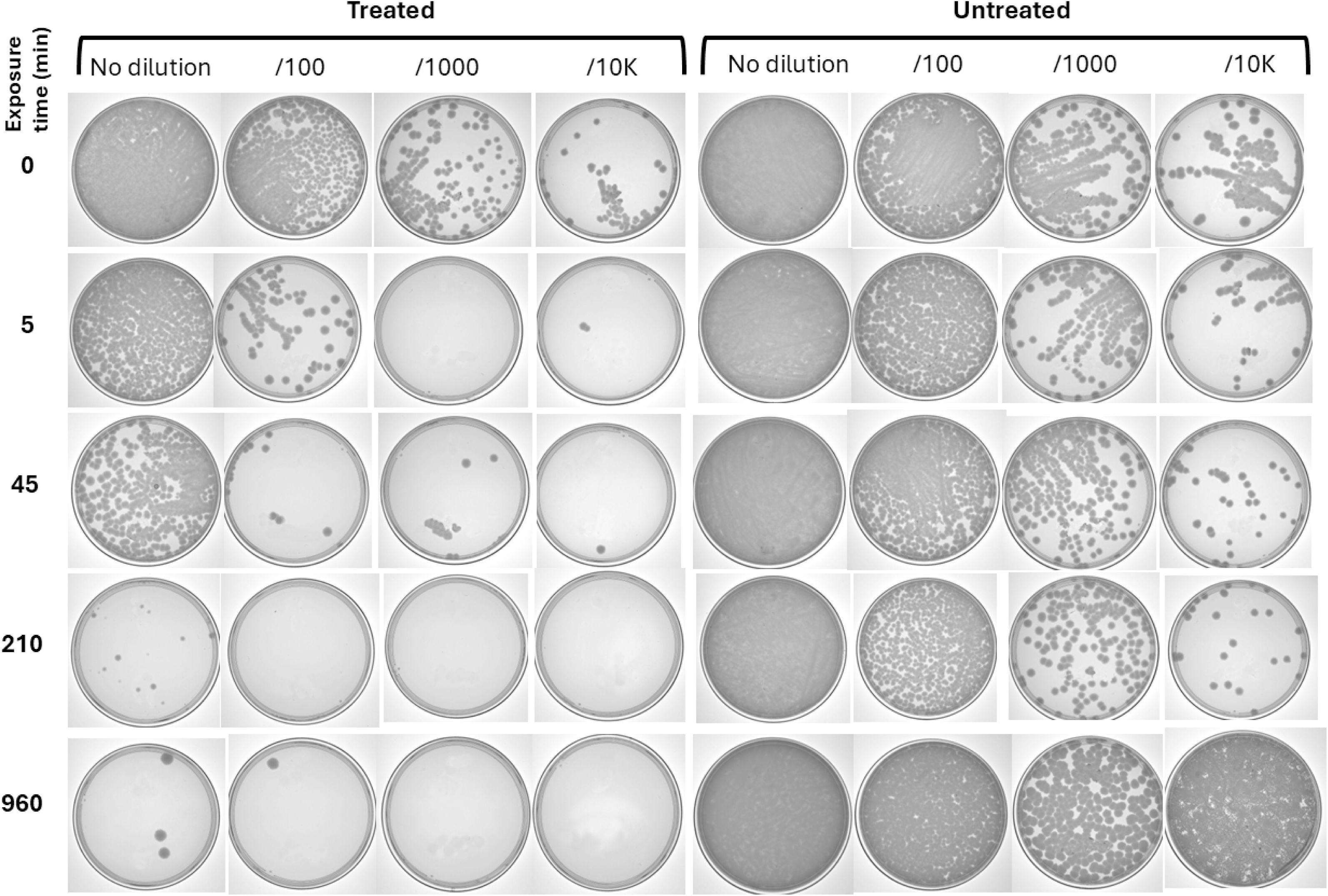
Prolonged exposure enhances efficacy of treatment- *B. cereus*. Exponentially growing cultures (10^7^ CFU/mL) of *B. cereus* bacteria were exposed for the indicated times to H₂O (right panels, untreated) or to 0.16% formic acid/0.4 mM CuSO₄. Serial dilutions (as indicated) were plated for CFU enumeration. Images are representative of at least three independent experiments.

**Supplementary Figure 4.**
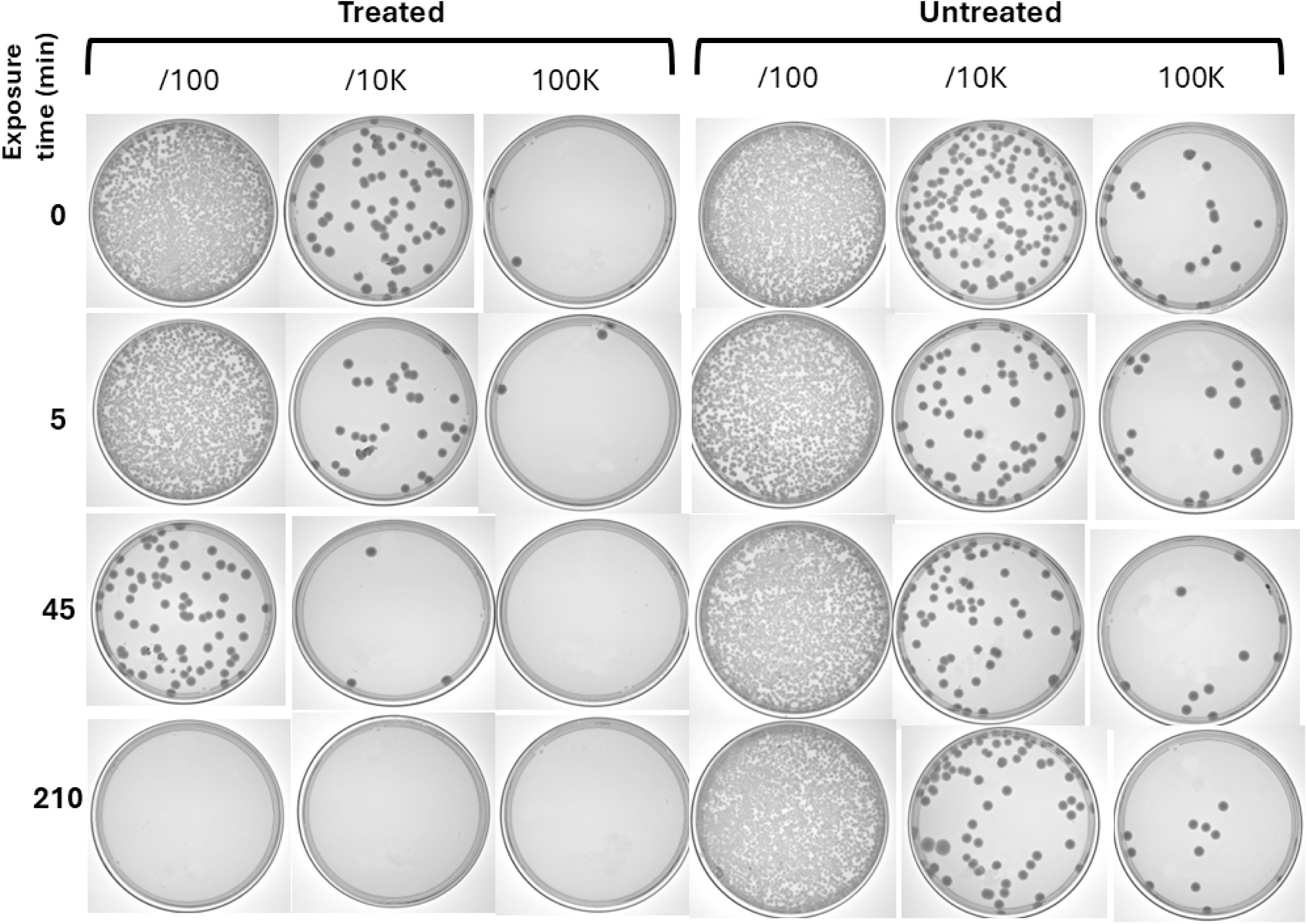
Prolonged exposure enhances efficacy of treatment- *P. aeruginosa*. Same as Supplementary Figure 3, only cultures of Pseudomonas aeruginosa were treated with 1% acetic acid/1 mM CuSO₄.

**Supplementary Figure 5.**
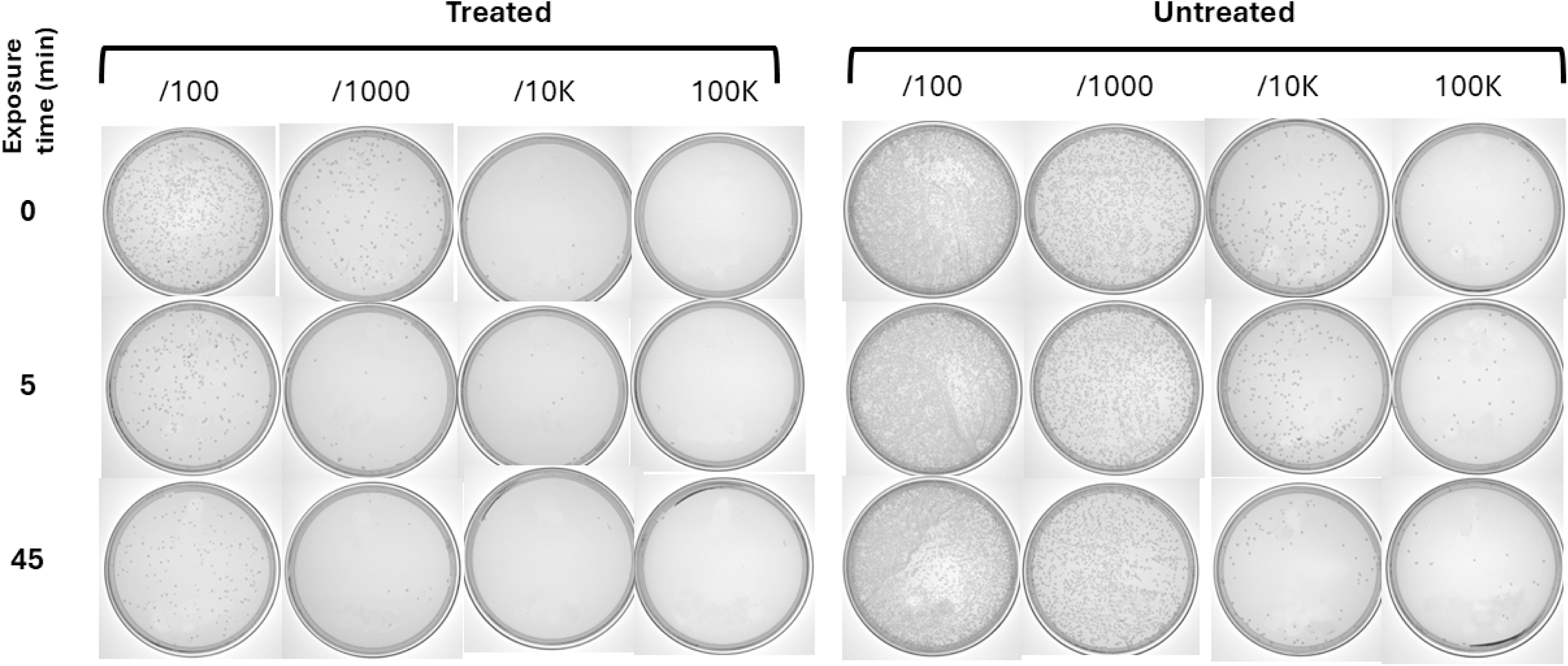
Prolonged exposure enhances efficacy of treatment- *Y. enterocolitica*. Same as Supplementary Figure 3, only cultures of *Y. enterocolitica* were treated with 0.36% formic acid/0.9 mM CuSO₄.

